# Cortical network mechanisms of response inhibition

**DOI:** 10.1101/2020.02.09.940841

**Authors:** Michael Schaum, Edoardo Pinzuti, Alexandra Sebastian, Klaus Lieb, Pascal Fries, Arian Mobascher, Patrick Jung, Michael Wibral, Oliver Tüscher

## Abstract

Both the right inferior frontal gyrus (rIFG) and the pre-supplementary motor area (pre-SMA) are crucial for successful response inhibition. However, the particular functional roles of those two regions have been controversially debated for more than a decade now. It is unclear whether the rIFG directly initiates stopping or serves an attentional function, whereas the stopping is triggered by the pre-SMA. The current multimodal MEG/fMRI study sought to clarify the role and temporal activation order of both regions in response inhibition using a selective stopping task. This task dissociates inhibitory from attentional processes. Our results reliably reveal a temporal precedence of rIFG over pre-SMA. Moreover, connectivity during response inhibition is directed from rIFG to pre-SMA and predicts stopping performance. Response inhibition is implemented via beta-band oscillations. Our findings support the hypothesis that response inhibition is initiated by the rIFG as a form of attention-independent top-down control.

## Introduction

Response inhibition is an essential component of cognitive control. Many neuropsychiatric disorders like attention deficit hyperactivity disorder (ADHD), obsessive-compulsive disorder (OCD), and Parkinson’s disease (PD) demonstrate the severe impact of impaired response inhibition (Schachar and Logan, 1990; De Wit et al., 2012; Voon and Dalley, 2011) on mental health. The classical stop-signal task (SST) has been used to operationalize response inhibition in many studies, thereby playing a major role in defining research on cognitive control (Verbruggen et al., 2019). In the go trials of this task, participants have to respond to a go signal by executing an immediate motor response, e.g. performing a button press. In less frequent stop trials, the go signal is followed by a stop signal after a variable delay and participants have to cancel the planned or ongoing motor response. A well-developed theoretical framework describing the cognitive processes underlying response inhibition relies on a so-called independent race model, which assumes the independence of go and stop processes (Logan et al., 1984; Verbruggen et al., 2019). Depending on which process finishes first, the stop process, which is initiated after the variable stop-signal delay, or the go process, which has been started already before, the response can be inhibited successfully or not. Based on this framework, the length of the stopping process can be estimated by subtracting the mean stop signal delay from the mean go RT. This length is also called stop signal reaction time (SSRT), which represents a behavioral measure for response inhibition in the absence of a motor reaction.

So far, numerous studies using the SST provided consistent evidence that response inhibition crucially depends on two cortical regions: the inferior frontal cortex (IFC) and the pre-supplementary motor area (pre-SMA) (Wessel and Aron, 2017). The particular functional roles of those two regions within response inhibition has been controversially debated for more than a decade now (Aron et al., 2004, 2014; Hampshire and Sharp, 2015a; Aron et al., 2015; Hampshire and Sharp, 2015b). While a range of studies on response inhibition point towards a single inhibitory module within the IFC that directly initiates stopping (Aron et al., 2003, 2014), other studies assign this role to pre-SMA (Li et al., 2006; Nachev et al., 2007). As an alternative, some authors suggest that the functional role of IFC within response inhibition is either the encoding of context-specific task rules or simply the attentional detection of the need for response inhibition, whereas its initiation is triggered by the pre-SMA (Duann et al., 2009; Sharp et al., 2010; Rae et al., 2015; Xu et al., 2017). A more critical view questions the behavioural construct of motor response inhibition in general. Hampshire and Sharp (2015a) suggest that there is no unitary response inhibition module and claim that the IFC is part of a domain-general control network (see also Erika-Florence et al., 2014). However, in the attempts to experimentally define the differential roles of IFC and pre-SMA, the temporal activation order of those regions has become a decisive, yet unresolved question in the field (Allen et al., 2018; Swann et al., 2012).

Two methodological issues may explain why this topic has so far remained unresolved. One is a lack of temporal resolution in many respective studies, the other the specific design of the standard SST. The majority of studies used fMRI, while only few studies in humans have addressed the temporal activation order of IFC and pre-SMA using techniques providing both, high spatial and high temporal resolution at the same time. One study used electrocorticography (ECoG) in a single patient and concluded that pre-SMA precedes the IFC activation (Swann et al., 2012). In contrast, two recent MEG studies suggest that both regions may be simultaneously active and that there is no temporal or functional primacy (Allen et al., 2018; Jha et al., 2015). However, the latter two studies may be limited by the design of the standard SST, which does not allow to disentangle attentional from inhibitory processes, and hence, to unequivocally define the functional roles of IFC and pre-SMA. To specifically control for the attentional load involved in the SST, Sharp et al. (2010) used a selective stopping task. This task comprises additionally attentional capture go trials that are identical to stop trials in terms of timing and frequency, but participants have to continue with their already prepared go response (Figure 1). Contrasting both conditions (stop vs. attentional capture go trials) not only allows to separate the two cognitive processes (attentional processing and response inhibition; Sebastian et al., 2016), but more importantly also enables direct temporal comparison of stop and go processes in terms of the onset of the related processes which is a prerequisite of high temporal resolution studies. So far, to our best knowledge, only three studies used a selective stopping task with an attentional capture go (“continue”) condition in connection with a high temporal resolution technique. Swann et al. (2012) used ECoG in a single patient, and Wessel et al. (2013) in four patients, respectively, but Wessel et al. (2013) could only cover the rIFC region, not pre-SMA. Sánchez-Carmona et al. (2016) used EEG, but did not analyze the temporal activation order of both regions. However, the majority of studies providing high temporal resolution relied on a standard SST (Allen et al., 2018; Bartoli et al., 2018; Fonken et al., 2016; Jha et al., 2015).

**Figure 1:**
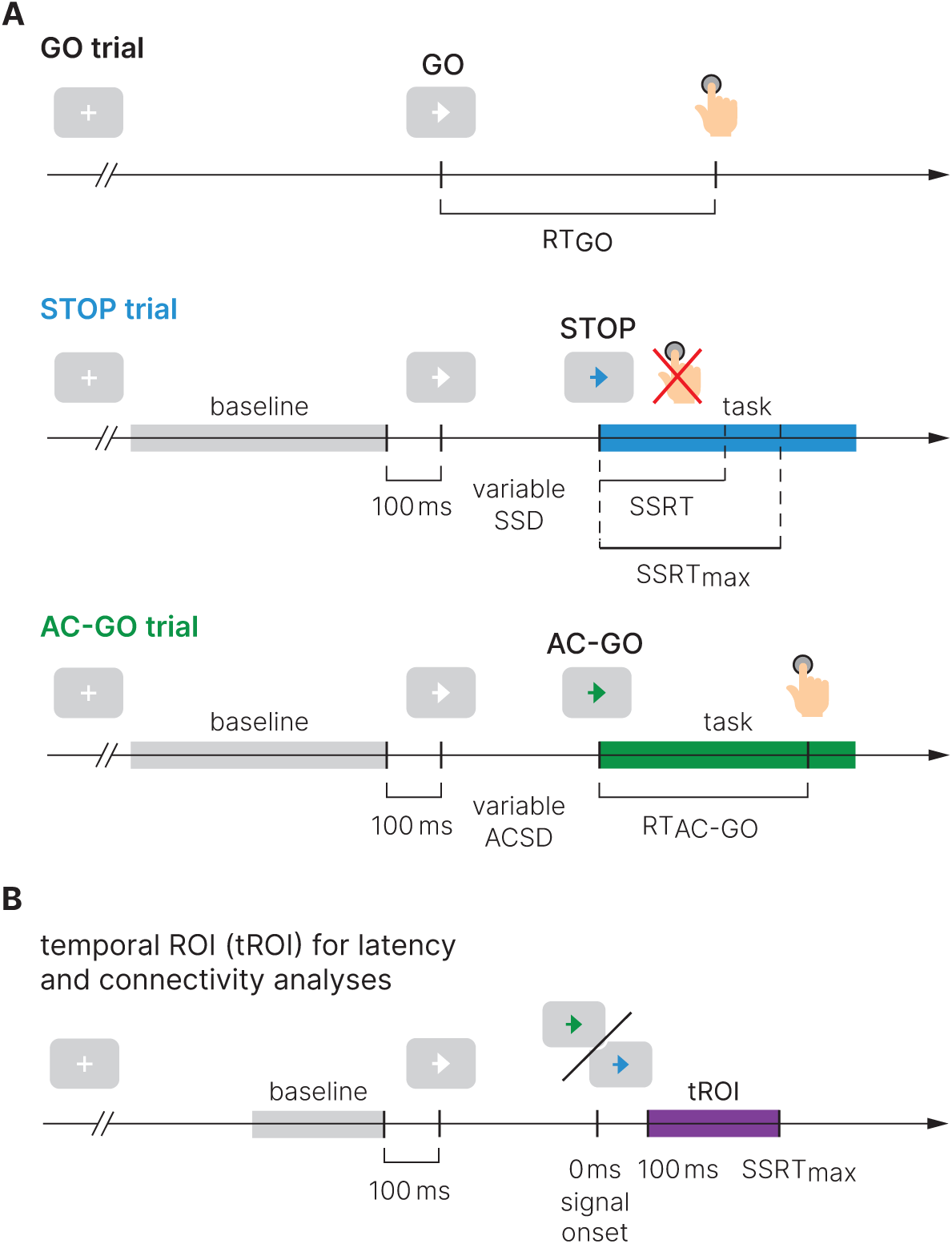
Experimental design of the selective stopping task and trial definition used in this study. **A**, The task comprised three conditions: a GO condition (50 % of all trials), a STOP condition (25 % of all trials), and an attentional capture GO (AC-GO) condition (25 % of all trials). The variable stop signal delay (SSD) was adapted to the participants’ performance to yield a probability of 50% of successful response inhibitions per block. Task segments of the trials were aligned to the STOP/AC-GO signal (stop trials in blue, AC-GO trials in green). Baseline segments (grey) are aligned to end 100 ms before GO signal onset. To reduce contamination with button-press related motor activity within the temporal region of interest, selective filtering was applied on AC-GO trials by excluding trials with RT_AC-GO_ < SSRT (median SSRT across participants, 237 ms). **B**, Temporal region of interest (violet) for latency and connectivity analysis starts at 100 ms, where early visual processing is likely complete and ends at SSRT_max_ (maximal SSRT across participants, 350 ms). *RT* reaction time, *SSRT* stop-signal reaction time, *SSD* stop-signal delay, *ACSD* attentional capture signal delay.

The current study aims to answer long-standing unresolved questions concerning the roles and timing of IFC and pre-SMA during response inhibition, i. e., which region initiates inhibitory control and which region exerts a putative causal influence on the other. We therefore used a selective stopping task in combination with high temporal resolution MEG recordings. To validate MEG source reconstruction we used the high spatial resolution of fMRI scans in a multimodal imaging approach. Based on previous findings, we hypothezised a predominant role of beta-band activity in both regions during the time of stopping (Kühn et al. (2004); Swann et al. (2009, 2012); Fonken et al. (2016)), and aimed to define the temporal activation order and connectivity of IFC and pre-SMA during response inhibition.

## Results

### Behavioral Results

Healthy participants (*n* = 62; three were excluded from further analysis, see Methods) performed a selective stopping task with an additional attentional control condition (Figure 1). Depending on different signal colors, participants were instructed to respond with a button press, to try to inhibit or to continue their already initiated go response. The mean stop signal delay (SSD) was 299.1 ± 132.6 ms (mean ± sd) and led to a probability of responding on a STOP trial close to 50 % (48.1 % ± 3.5 %) proving the adherence of subjects to the task rules and the successful operation of the staircase procedure. While for successful STOP trials (sSTOP) the mean SSD was 288.2 ± 124.7 ms, it was 312.7 ± 128.8 ms for unsuccessful STOP (uSTOP) trials. The mean attentional capture signal delay (ACSD) for correct attentional capture GO (cAC-GO) trials was 302.0 ± 124.2 ms. A repeated-measures ANOVA based on RT revealed a main effect of condition (*F* = 467.1, *p* < 0.001). Bonferroni-corrected post hoc comparisons revealed that mean RT in cAC-GO trials (588.2 ± 13.5 ms, group mean ± standard error of the mean) was significantly longer as compared to GO trials (550.1 ± 14.0 ms) and uSTOP trials (503.7 ± 13.1 ms). Mean RT in uSTOP trials was significantly shorter as compared to GO trials (*p* < 0.001 for all post hoc tests; Figure 2). Participants performed accurately as indicated by low omission error rates in GO (2.3 ± 2.8 %, mean ± sd) and AC-GO trials (1.2 ± 1.4 %). All these behavioral results of the MEG study are in line with the ones of the fMRI study (Sebastian et al., 2016). SSRT was calculated by the mean and integration method (see Methods). While the integration method tends to underestimate SSRT, the mean method tends to overestimate it (Verbruggen et al., 2013). Therefore we averaged the values of both methods as suggested by Jha et al. (2015), resulting in a median SSRT of 237.4 ± 39.2 ms. The maximal SSRT across participants (SSRT_max_), based on these averaged SSRT values, was 349.9 ms.

**Figure 2:**
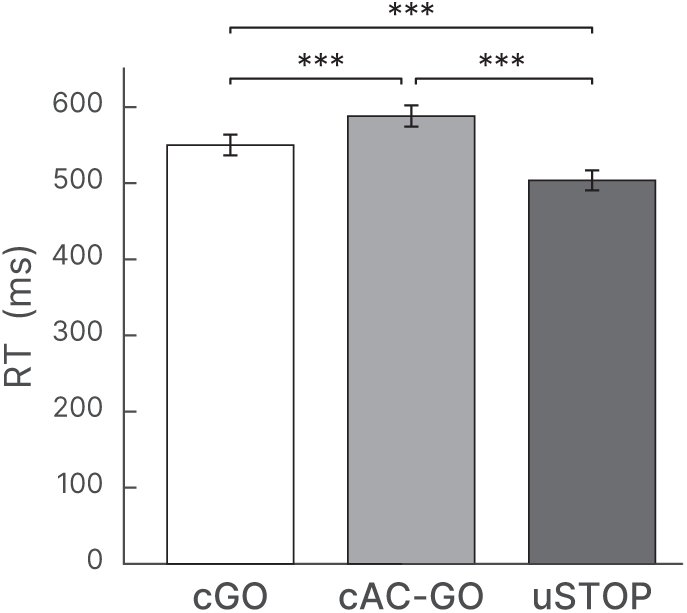
Behavioral results for the stop signal task (mean ± standard error of the mean). Reaction time (RT) in correct GO (cGO), correct attentional capture GO (cAC-GO), and unsuccessful stop trials (uSTOP). *** *p* < 0.001 based on Bonferroni-corrected post hoc comparisons of a repeated-measures ANOVA.

To exclude an effect of signal color, the experiment was designed in a cross-balanced manner, i. e., the attribution of signal color for stopping (blue/green) to trial type (STOP/AC-GO) was balanced across subjects. A two-sample *t*-test with signal color as between factor revealed that the stopping latency as measured by the SSRT did not differ significantly between groups (*t* = 0.376, *p* = 0.708). A mixed-design ANOVA with signal color as between factor and reaction time (RT; GO vs. AC-GO) as within factor further revealed no influence of attribution of color to trial type, as no interaction of these two factors was present (F = 1.745, p = 0.192).

### Source activation

To analyze the dynamics and connectivity of response inhibition, we first identified a set of cortical sources involved in successful stopping. Importantly, we aimed to identify sources that are actually active in the acquired data and not only motivated by literature (Gross et al., 2013). First, to find appropriate parameters for source reconstruction using a beamformer method, the spectral power at sensor level was analyzed for both conditions pooled, sSTOP and cAC-GO trials. We a-priori defined a temporal region of interest (tROI) starting after the presumed completion of early visual processing and ending with SSRT_max_ (100 to 350 ms, see Methods). We compared spectral power between tROI and an equally long baseline epoch (see Supplemental Information, Figure S1). The cluster-based permutation test revealed one cluster with significant power decrease (12.0 to 31.9 Hz, beta band) and one cluster with significant power increase (63.8 to 87.7 Hz, gamma band). Second, to reveal effects related to response inhibition, the reconstructed source power during the tROI was contrasted between both conditions (sSTOP and cAC-GO) for each of those two frequency bands. The contrast revealed that beta-band power was increased in sSTOP compared to cAC-GO trials in a network of pre-motor and pre-frontal areas (Figure 3A). One main cluster of increased beta-band power was located in the right inferior frontal cortex with its maximum in the rIFG pars opercularis. Increased beta-band power was also present in the pre-SMA, l-MFG, and bilateral premotor regions (Table 1, Figure 3A). The same analysis performed for the gamma band did not reveal a significant cluster. To further validate the MEG source reconstruction, a subset of participants in the MEG experiment (*n* = 31) plus 45 additional participants were recorded using fMRI (resulting in *n* = 76). Here, the contrast sSTOP > AC-GO revealed activation in two rIFG subregions (p. triangularis, BA45, *x* = 52, *y* = 22, *z* = 4, *t* = 10.96; p. opercularis, BA44, *x* = 52, *y* = 18, *z* = 14, *t* = 9.42) and pre-SMA (*x* = 16, *y* = 14, *z* = 64, *t* = 9.45) as sub-maxima in the biggest cluster obtained (13438 voxels, see Figure 3B). A smaller cluster (85 voxels) also showed pre-SMA related activation (*x* = –14, *y* = 18, *z* = 66, *t* = 5.63). The peak coordinates obtained by the MEG source construction in the beta band were closely located to the peak voxels identified fMRI, i. e., the minimal distance between MEG and fMRI sources was 6.6 mm for the rIFG areas and 15.4 mm for the pre-SMA areas.

**Table 1:** MEG source reconstruction in the beta band. MNI coordinates obtained as peak voxels when contrasting sSTOP versus cAC-GO trials.

| Region | MNI [mm] |  |  | *t*-value | Abbr. |
| --- | --- | --- | --- | --- | --- |
|  | *x* | *y* | *z* |  |  |
| Inferior Frontal Gyrus BA44/45 R | 50 | 20 | 20 | 3.654 | rIFG |
| Inferior Frontal Gyrus BA44/45 L | -50 | 10 | 20 | 2.820 | l-IFG |
| Anterior Insula / left IFG | -40 | 30 | 0 | 2.894 | lAIns |
| Pre-supplemental motor area | 0 | 20 | 60 | 3.878 | pre-SMA |
| Middle Frontal Gyrus | -40 | 0 | 50 | 3.360 | lMFG |
| Premotor cortex BA6 R | 10 | -10 | 50 | 4.015 | rPMC |
| Premotor cortex BA6 L | -20 | -10 | 60 | 3.571 | lPMC |
| Fornix | 0 | 0 | 10 | 3.740 | Fornix |

**Figure 3:**
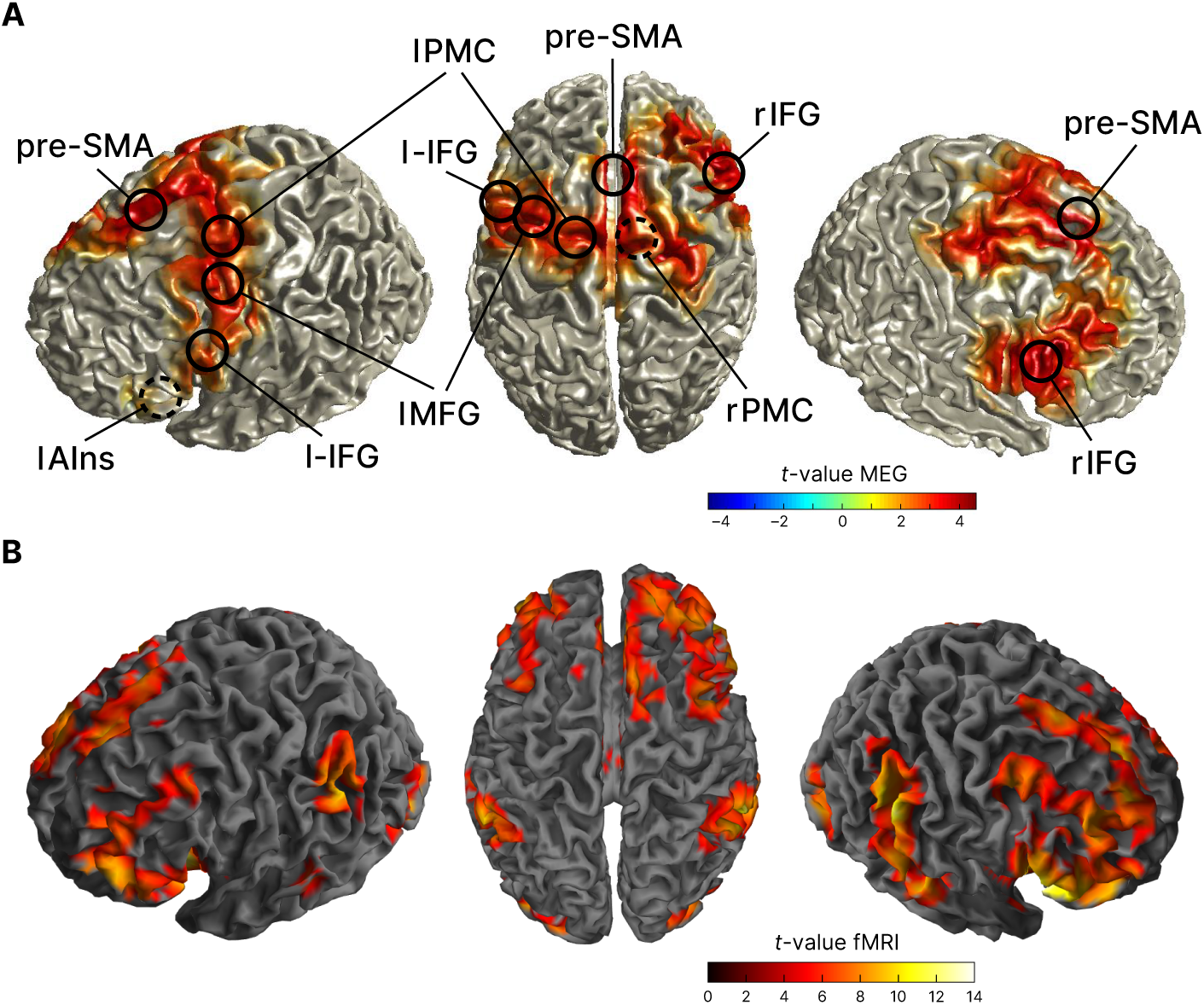
Sources involved in successful stopping. **A**, Source reconstruction of MEG data in the beta band (12–32 Hz, 100–373 ms after STOP/AC-GO signal onset). Surface plots show *t*-values of significant clusters when contrasting sSTOP and cAC-GO trials (*α*_cluster_ = 0.05, two-tailed test). Peak voxels (local extrema) of these clusters are highlighted and labelled (MNI coordinates are shown in Table 1). **B**, fMRI activation maps for the contrasts sSTOP > AC-GO; map thresholded *p*_FWE_ < 0.05, cluster extent *k* = 5 voxels). Data are taken from Sebastian et al. (2017).

### Temporal precedence of rIFG over pre-SMA activation

To test our main hypothesis concerning the initiation of response inhibition, we analyzed the temporal development of beta-band power across the sources obtained by contrasting *z*-transformed sSTOP with cAC-GO trials. The rIFG was the source that showed the earliest beta-band power difference compared to all other sources (Figure 4A). According to our hypothesis, we tested if the beta-band power difference occurred earlier or later at the rIFG than at the pre-SMA. Since cluster-based permutation tests do not establish significance of latency differences between conditions (Sassenhagen and Draschkow, 2019), we tested the temporal lead of rIFG activity onset by subtracting *z*-transformed cAC-GO trials from sSTOP trials (Figure 4B, group level). Onsets were defined by using a percentage threshold of the range between zero power and the first local power maximum found in the tROI (Figure 4C). These onsets were then used as input for a permutation test that revealed a significant latency onset difference between both sources (*p* = 0.0240, two-tailed test, mean onset latency rIFG: 137 ± 46 ms, and pre-SMA: 159 ± 52ms. This difference refers to a 25 % onset threshold, but it was significant for 10 %, 30 %, and 50 % thresholds, too. For this test, seven subjects had to be excluded because no positive peak could be found within the tROI.

**Figure 4:**
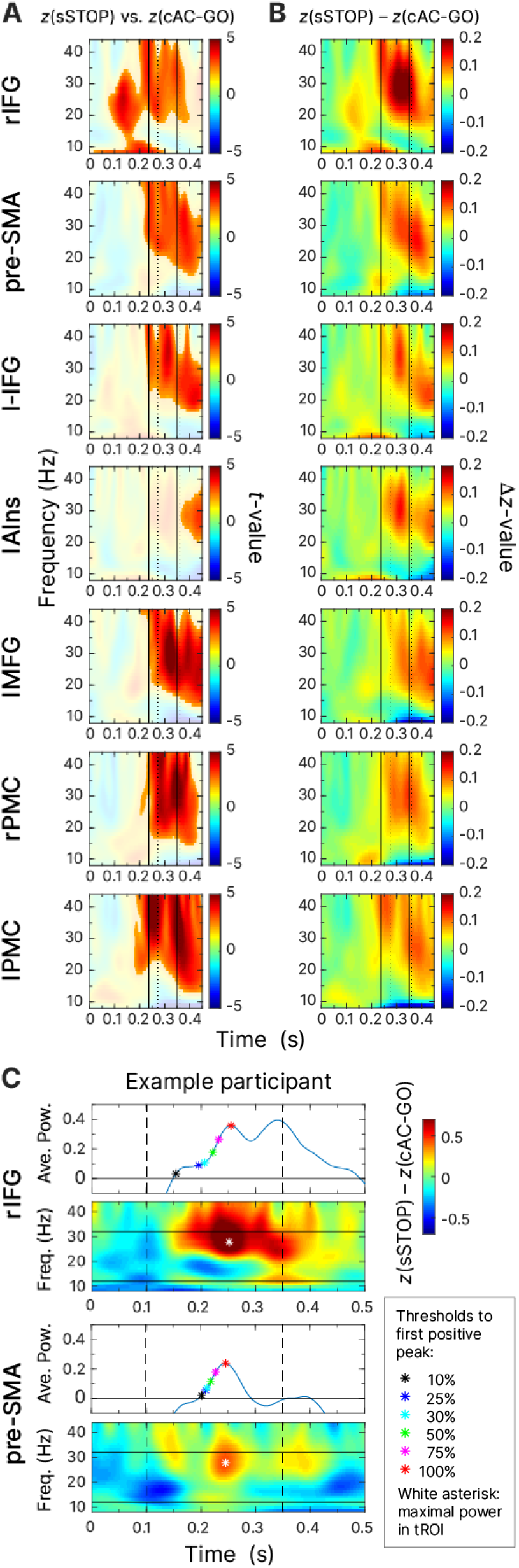
**A and B**, Beta-band time-frequency representations (TFRs) of virtual channels placed at sources identified. First and second solid line indicate SSRT and SSRT_max_, first and second dashed line indicate 10% and 50%-percentiles of RT_AC-GO_ for selected trials with RT_AC-GO_ > SSRT. **A**, contrast of *z*-transformed sSTOP versus cAC-GO trials (values inside the significant clusters are displayed opaque). **B**, absolute difference between *z*-transformed sSTOP and cAC-GO trials averaged across subjects. **C**, TFRs of this *z*-transformed difference for an example participant. In the TFR plots, dashed lines represent the tROI (100–350 ms) and solid lines limit the frequency band used for onset latency analysis. The white asterisk highlights the maximal power difference found in this time-frequency window. The subplots above the TFRs show power values averaged over frequencies (12–32 Hz). This curve was used to define the onset latency, e. the time point at which the source power exceeded a threshold of 10, 25, 30, 50, 75, and 100 % of the range between zero power and the first power maximum found in the tROI.

### Beta-band source power correlates with inhibitory perfomance

On the basis of the remaining participants, we correlated source power with SSRT to test behavioral significance of beta-band (12–32 Hz) power differences during the tROI. To this end, power differences between *z*-transformed sSTOP and cAC-GO trials were averaged over trials for each participant at the peak voxel of rIFG and pre-SMA sources according to our hypothesis. Each of both peak voxels was identified as local extremum within a signficant cluster at group level (see Methods). The maximal power difference found in the beta band within the tROI was negatively correlated with SSRT values for the rIFG (*ρ* = –0.246, *p* = 0.043, Spearman, one-sided test), but not for pre-SMA (*ρ* = –0.121, *p* = 0.200, Figure 5; two outliers above three standard deviations were excluded from both regressions, *n* = 50). t

**Figure 5:**
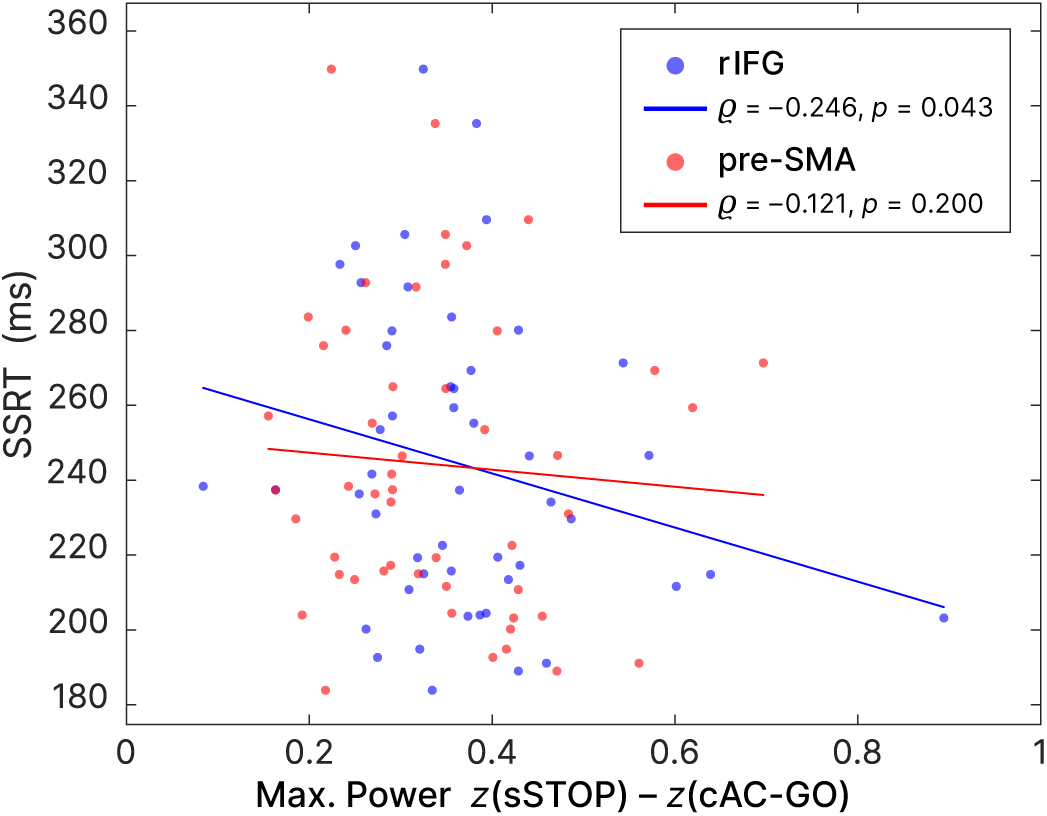
The maximum (*z*-transformed) power found in the beta band (12–32 Hz) within the tROI (100–350 ms) was negatively correlated across subjects with SSRT values for rIFG, but not for preSMA.

To further ensure that the earlier onset latency of rIFG compared to pre-SMA does not rely on the specific time-frequency transformation method used and the parameters applied for onset latency definition, we used an orthogonal approach to define the onset latency of both regions. Here, we used broadband signal time courses instead of a frequency-specific analysis. We performed a time-resolved support-vector-machine (SVM) analysis in order to classify sSTOP versus cAC-GO trials and to identify the discriminability onset between both conditions, separately for rIFG and pre-SMA. This SVM-based approach revealed a robust above-chance classification starting from 137 ms (*p* = 0.0001) after STOP/AC-GO signal onset for the rIFG source and from 157 ms (*p* = 0.0001) for the pre-SMA source (Figure 6). To statistically compare individual classification onsets for the two sources, we performed an onset analysis of the final classification time series. This analysis showed a significantly earlier rIFG onset (mean onset 185 ms) compared to pre-SMA (mean onset 207 ms) at a threshold of 25 % (*p* = 0.002; 13 subjects were excluded because they did not meet inclusion criteria, see Methods). This result was not biased by the selected threshold since all other thresholds gave similar results. The SVM results indicate that the rIFG source contained information to distinguish sSTOP and cAC-GO trials at an earlier time point than the pre-SMA. Overall, the analysis of time domain data corroborates the significantly earlier activation of rIFG in the beta-band power analysis, suggesting that this area plays a leading role in initiating response inhibition.

**Figure 6:**
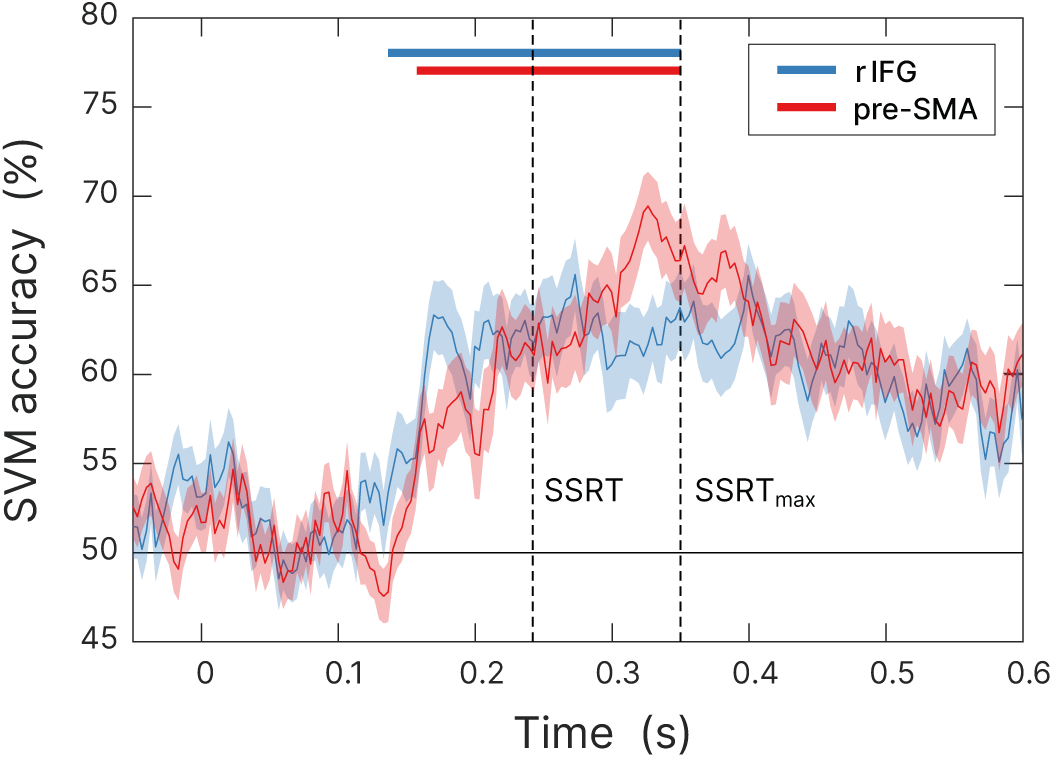
Time-resolved support-vector-machine (SVM) analysis classified both trial types robustly above chance within the tROI (100–350 ms) across subjects. Robust above-chance classification was tested between 0–350 ms. Significant time windows within this range for rIFG and pre-SMA are highlighted by colored bars. A statistical test for individual differences in onset latency between both regions revealed that sSTOP and cAC-GO trials could be discriminated 22 ms earlier in rIFG (mean onset 185 ms) than in pre-SMA (mean onset 207 ms) at a 25 % threshold.

### rIFG is the dominant sender of information during response inhibition

To further characterize the functional role of rIFG and pre-SMA during response inhibition, we analyzed their connectivity pattern using nonparametric conditional Granger causality (cGC). We contrasted spectral cGC of sSTOP and cAC-GO trials in the tROI. Therefore, we computed the cGC from rIFG to pre-SMA and vice versa, conditional on all other five active cortical sources as obtained by the source reconstruction contrast. The cluster-based permutation test (see Methods) revealed statistical differences between sSTOP and cAC-GO conditions in the direction rIFG to pre-SMA within the (high) beta band (*p*_cluster_ = 0.0074, corrected for the two links tested), but not vice-versa (Figure 7A).

**Figure 7:**
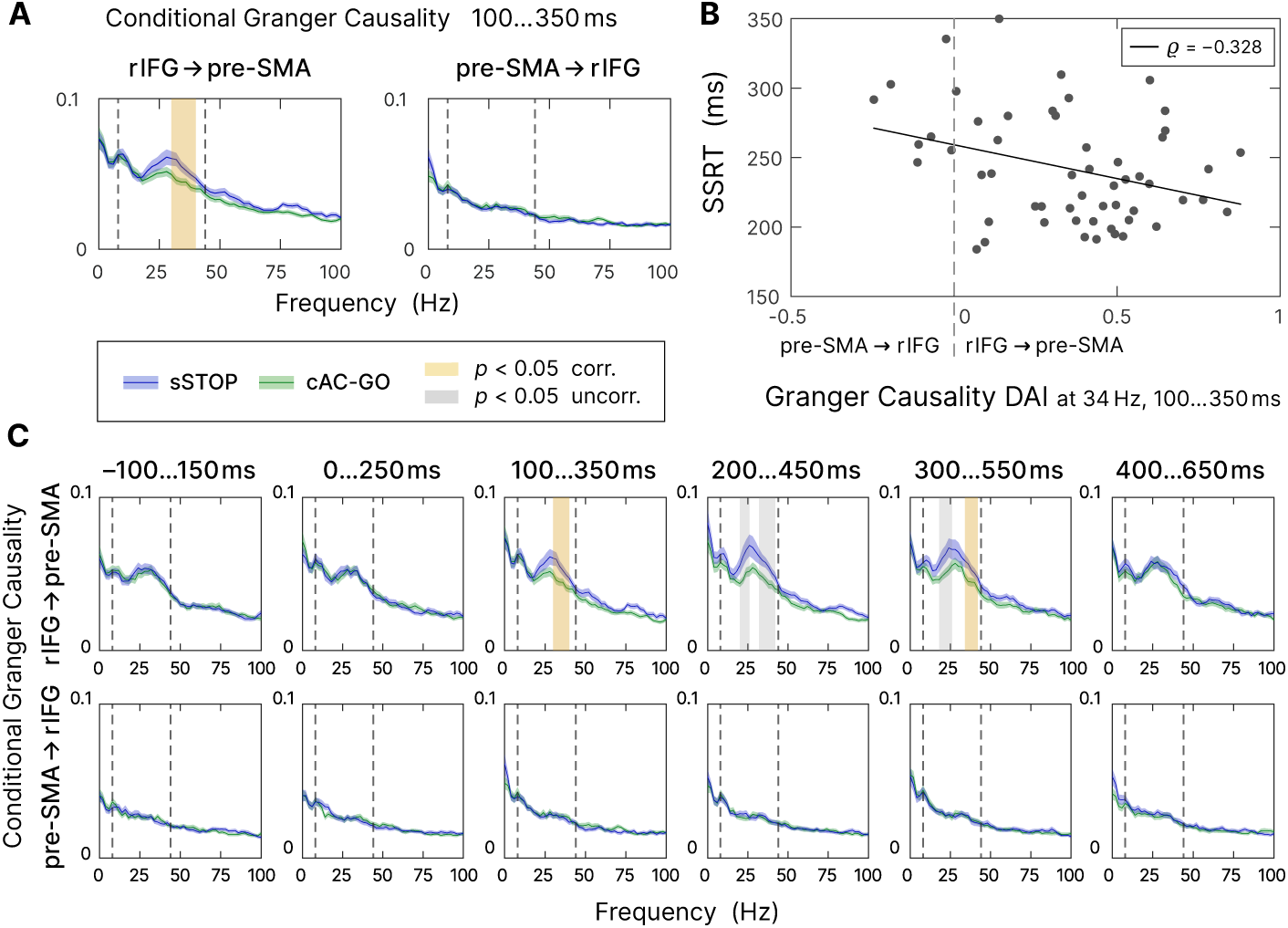
Connectivity analysis. **A**, Spectral-resolved conditional Granger causality (cGC) between rIFG and pre-SMA. Blue curve: cGC for sSTOP trials, green curve: cGC for cAC-GO trials, bounded lines: standard error of the mean. Significant differences between both conditions are highlighted in yellow if corrected for multiple comparisons, i. e., the two links tested, and in grey if uncorrected. The frequency range tested (8–44 Hz) is indicated by dashed lines. **B**, SSRT correlated with directionality asymmetry index (DAI) for rIFG and pre-SMA cGC values (sSTOP trials, cGC at 34 Hz in the tROI, 100 to 350 ms). Positive DAI corresponds to links from rIFG to pre-SMA, while negative DAI corresponds to links from pre-SMA to rIFG. The negative correlation indicates that participants with higher cGC from rIFG to pre-SMA are the better inhibitors (faster SSRT). **C**, Temporal evolution of cGC was analyzed post hoc between rIFG to pre-SMA (top) and pre-SMA to rIFG (bottom). A sliding window was shifted with 100 ms steps around the tROI. cGC from rIFG to pre-SMA (but not vice-versa) was significantly higher for sSTOP compared to cAC-GO trials in the tROI itself and in a later time window tested. Here, we corrected for testing six time windows and for testing in two directions (rIFG to pre-SMA and vice-versa). Same formatting as in **A**.

To assess if the connectivity between rIFG and pre-SMA was correlated with SSRT, we used the directed influence asymmetry index (DAI; Bastos et al., 2015) that captures the predominant direction of cGC. Normalization allows comparison of DAIs across frequencies (Michalareas et al., 2016). We correlated the DAI of the sSTOP condition with SSRT in the significant frequency range identified (in steps of 2 Hz, in the range from 30 to 40 Hz, resulting in six discrete frequencies; Figure 7A). Using a bootstrap method (Bastos et al., 2015), we found a significant negative correlation between DAI and SSRT at 34 Hz (*r* = –0.328, within a 99.17% confidence interval [–0.573; –0.030]). Hence, the better subjects were able to inhibit their response, i. e., the faster their SSRT was, the higher, positive DAI values they had, indicating a stronger connectivity from rIFG to pre-SMA, whereas slower inhibiting subjects showed the reverse pattern (negative DAI, stronger connectivity from pre-SMA to rIFG, Figure 7B). This demonstrated that the dominant direction of communication between rIFG and pre-SMA affects the stopping process significantly.

Finally, to test whether rIFG to pre-SMA connectivity was specific to the tROI of response inhibition, we tested the temporal evolution of the cGC contrast between sSTOP and cAC-GO trials. Therefore, we performed a sliding window analysis, moving backward and forward from the main tROI (100 to 350 ms) in steps of 100 ms (Figure 7C). This post hoc analysis revealed a decrease in rIFG to pre-SMA connectivity from 200 to 450 ms. Only the flanks of the beta band showed a difference in cGC between conditions, but this effect was not significant after the correction for multiple comparisons of six time windows times and of each direction tested (*p*(lower cluster) = 0.0052, *p*(upper cluster) = 0.0207, both uncorrected). However, in a later window (300 to 550 ms), the upper flank of this band showed a significant difference between conditions (*p*(upper cluster) = 0.0384, corrected, and *p*(lower cluster) = 0.0064, uncorrected). No significant differences were found in any time window before the tROI and after 550 ms, nor in the direction of pre-SMA to rIFG. These findings suggest that response inhibition is initiated within a crucial time range by specific rIFG to pre-SMA connectivity, the strength of which correlates with behavioral performance.

To exclude that these results were biased by the cAC-GO trial selection (only including trials with RT_AC-GO_ > SSRT), we performed all analyses on randomly selected cAC-GO trials. Overall, these results were comparable to the reported ones.

## Discussion

The neural network mechanisms of response inhibition are central to the understanding of the neurobiology of cognitive control and have been the matter of a long-standing debate. Here, we tested which of the two main regions proposed as the cortical initiator of response inhibition, rIFG and pre-SMA, respectively, is activated first and which exerts a putative causal influence on the other.

Our study, using the high temporal resolution of MEG and a relatively large cohort of subjects, is the first to show a significantly earlier activation in the rIFG compared to the pre-SMA. In particular, rIFG beta-band activity precedes pre-SMA activity by approximately 22 ms – as independently verified by a latency and a time-resolved SVM analysis. Connectivity analysis showed that rIFG sends information in the beta band for successful stopping to pre-SMA but not vice versa. The behavioral significance of beta-band power as well as connectivity from rIFG to pre-SMA in the beta band was demonstrated by their correlation with stopping performance.

### rIFG initiates response inhibition

So far, the temporal activation order of rIFG and pre-SMA in response inhibition has been investigated at source level with high temporal resolution (MEG or ECoG) by only a limited number of previous studies using a stop signal task (Allen et al., 2018; Jha et al., 2015; Swann et al., 2012). Allen et al. (2018) and Jha et al. (2015) both found that the pre-SMA and rIFG were simultaneously activated in the time window between STOP signal and SSRT. In contrast to these MEG studies, one ECoG study in a single patient concluded that pre-SMA precedes the IFC activation (Swann et al., 2012). There may be three reasons for these diverse findings. First, these previous studies rely on a comparably small sample sizes (*n* = 19, *n* = 9, *n* = 1, respectively) compared to the current study (*n* = 59). A second reason is the use of different SST paradigms, i. e. non-selective (standard) SST (Allen et al., 2018; Jha et al., 2015) versus a selective stopping task (Swann et al., 2012, and current study). Contrasting sSTOP trials with uSTOP or GO trials in non-selective SSTs entails several issues (see discussion below, *Selective stopping* task suggests that rIFG does not serve an attentional role). A third reason is that results may depend on the way onset latencies are defined (see discussion of limitations by Bartoli et al., 2018). We therefore validated our findings based on time-frequency representations (TFR) by a second, orthogonal approach. Here, sSTOP and cAC-GO trials were classified using a time-resolved SVM. The differences between sSTOP and cAC-GO trials could be detected 22 ms earlier in rIFG than in pre-SMA in the SVM analysis, independently replicating our result on latency differences in the TFR analysis. Since the temporal onset of discriminability between conditions was found already before the SSRT, this suggests that rIFG activity is indeed pivotal for initiating stopping.

### Beta-band power in rIFG relates to inhibitory performance

We further hypothesized that if activity in the rIFG is a neural correlate for response inhibition, there should be a correlation with inhibitory performance at behavioral level, i. e., SSRT. Indeed, the power maximum in the beta band correlated negatively with SSRT in the rIFG only, but not in the pre-SMA. This is in line with several fMRI studies (Aron, 2006; Aron et al., 2007; Rubia et al., 2007), suggesting that subjects with greater activation in rIFG inhibited more quickly (shorter SSRT). This finding additionally supports the dominant role of the rIFG for initiating stopping.

In addition, we sought to clarify if the temporal precedence of rIFG over pre-SMA and the correlation of beta-band power in rIFG with SSRT depend on the outcome of the stopping process (successful vs. unsuccessful stop). Therefore, both analyses were applied on the contrast between uSTOP and cAC-GO trials, instead of the contrast between sSTOP and cAC-GO trials. The temporal activation pattern, i. e. rIFG before pre-SMA, was nearly the same for both contrasts (see Supplemental Information, *Exploratory analysis of uSTOP trials*). However, the correlation of beta-band power with SSRT was only significant for the difference between sSTOP and cAC-GO trials (Figure S2). Hence, the temporal precedence of rIFG over pre-SMA may indeed be relevant for stopping in general, while at the same time, the degree of control by the rIFG is responsible for the success and the performance of stopping.

### Dominance of rIFG during response inhibition supports its role as controller

To further elucidate the role of rIFG in response inhibition, we analyzed the connectivity between rIFG and pre-SMA using spectrally-resolved conditional Granger causality analysis (cGCA). This method measures directed information transfer between two regions, i. e., it quantifies predictive relationships within the network (Granger, 1969; Geweke, 1982). If the rIFG indeed has the function of a top-down controller of response inhibition, then it should exert an ongoing causal influence on the pre-SMA across the time-window of response inhibition. Indeed, a significant difference between sSTOP and cAC-GO in cGC influence from rIFG to pre-SMA was detected, but not vice-versa. In agreement with our hypothesis, significant results were found within the beta band in particular. The observed directionality is in line with cGCA in a comparably high-powered fMRI study (*n* = 57, Duann et al., 2009), yet inconsistent with cGCA of a somewhat lower powered MEG study (*n* = 19, Allen et al., 2018). Even more, the strength of this directional asymmetry (DAI) was correlated to stopping performance as measured by SSRT. In addition, a DCM analysis of an EEG study available in preprint (Fine et al., 2019) showed that inhibitory connectivity from rIFG to pre-SMA during STOP trials was also negatively related to SSRT when transcranial focused ultrasound (tFUS) was applied. These findings suggest that connectivity directed from rIFG to pre-SMA per se reflects a state in which rIFG is prepared to send a signal to pre-SMA to execute stopping, while the strength of this directionality influences the performance of stopping. To test whether the directionality from rIFG to pre-SMA was specific for response inhibition, or just related to other processes, we analyzed connectivity between rIFG and pre-SMA before and after the (a priori defined) temporal ROI (tROI, 100–350 ms) using a sliding window approach. Thus, we could confirm that beta-band related connectivity between rIFG and pre-SMA was specific to the tROI and a subsequent time window, particularly only in the direction from rIFG to pre-SMA. In sum, the results of the spectrally resolved cGCA further support the leading role of rIFG in response inhibition, and the importance of beta-band oscillations in this process. Since connectivity from rIFG to pre-SMA was specifically established during the tROI and correlated with inhibitory performance, rIFG may be characterized as a brake in response inhibition.

### Inhibitory role of beta-band oscillations

Our results strongly support the interpretation that the rIFG acts as the initiator of response inhibition and that beta-band oscillations play an important role for its neural implementation. In general, beta-band oscillations are thought to be related to the maintenance of the current sensorimotor or cognitive state as hypothesized by a prominent model (Engel and Fries, 2010). In contrast, desynchronization of beta-band oscillations enables the transition into an active processing state (Miller et al., 2012), e. g. pressing a button. Our TFR results can be explained within the framework of this model, i. e., beta-band oscillations were less desynchronized in sSTOP trials than in cAC-GO trials, resulting in a net beta-band power increase (see Figure 4B). As soon as the stopping or motor response, respectively, is completed, beta-band oscillations start to resynchronize (compare Fonken et al., 2016). Notably, beta-band power in our study changed particularly before SSRT, which is in line with other studies (Wessel et al., 2013; Castiglione et al., 2019) and supports its inhibitory role.

### Selective stopping task suggests that rIFG does not serve an attentional role

A limitation of most previous neuroimaging studies on response inhibition is that sSTOP trials are usually contrasted with GO or uSTOP trials to identify inhibitory processes, while we contrasted sSTOP with cAC-GO trials. All three contrasts are indeed capable of removing basic perceptual activity due the presentation of the stimuli (Aron, 2006; Sharp et al., 2010). However, contrasting with uSTOP trials in turn involves error processing (Rubia et al., 2003) and an early response in motor cortices that has the potential to obscure activity related to the adjacent pre-SMA (Allen et al., 2018; Jerbi et al., 2007). Additionally, this contrast was criticized as overly conservative, because it assumes that inhibitory processes are also present in uSTOP trials (Boehler et al., 2010). Contrasting sSTOP with GO trials instead also introduces two issues: First, a less frequent event like the STOP signal is perceptually more salient and causes significant brain activation that is not directly based on inhibitory processing alone (Sharp et al., 2010). Second, STOP signals are delayed compared to the GO signals (i. e., SSD), hence directly contrasting both trial types entails the comparison of different time points within the stopping process (Figure 1). This is particularly important when high temporal resolution methods are used. Additionally, attentional and discriminative processing of the STOP signal are presumed to disturb the timing of go and stop processes compared to GO trials (Sharp et al., 2010). Both issues can be resolved by contrasting sSTOP with cAC-GO trials. First, we matched frequency and stimulus properties of sSTOP and cAC-GO signals. For STOP and AC-GO signals, blue and green arrows were used, respectively, and balanced across subjects. And indeed, the signal color had no effect on SSRT or with respect to RTs, on trial type (GO vs. AC-GO). Second, our behavioral data confirm the predicted increase in RTs after the appearance of an AC-GO signal compared to simple GO trials. There is independent evidence that the prolonged time required for the perceptual processing of the AC-GO signal is approximately the same as for the STOP signal compared to GO trials (Xu et al., 2017). The authors demonstrated at least the behavioral similarity of both conditions by comparing SSRT and “continue signal reaction time” (CSRT, i. e. approximately, the time between AC-GO signal and response, a measure adopted from Mayse et al., 2014). Hence, the contrast sSTOP vs. cAC-GO is the most appropriate to identify the neural correlates of response inhibition, independent of attentional and error processing (Sánchez-Carmona et al., 2016). The dissociation of inhibitory from attentional processes by an appropriate task design suggests that the rIFG does not, at least not primarily, serve an attentional role in response inhibition. However, although the rIFG has often been regarded as candidate for an inhibitory module and our results showed that response inhibition is indeed initiated by the rIFG, the rIFG might still be part of a more domain general cognitive control mechanism, implemented by frontoparietal networks (Erika-Florence et al., 2014; Hampshire and Sharp, 2015a).

### Limitations

One limitation of this study is that we could not analyze activity of the subthalamic nucleus (STN) which is an important subcortical node within the response inhibition network because of its direct relation to IFG (Aron et al., 2007; Duann et al., 2009; Jahfari et al., 2011; Rae et al., 2015; Xu et al., 2017). No STN activity was detected by our source reconstruction contrast, most likely because MEG is less sensitive to subcortical compared to cortical sources (Gross, 2019). Nonetheless, several studies with intracranial recordings detected beta-band oscillations within the STN (Kühn et al., 2004; Ray et al., 2012; Zavala et al., 2018; Fischer et al., 2018), also supporting the importance of beta-band oscillations in response inhibition.

### Conclusion

Our findings corroborate the hypothesis that response inhibition is initiated by the rIFG and implemented as a top-down control via synchronized activity in the beta frequency range (Aron et al., 2014). This brake is turned on in a time window around 100 ms to 200 ms after the STOP signal. Within the framework of the race model, rIFG may promote stopping processes and then signals to pre-SMA to proceed and execute the stopping process. Due to our task design, i. e., contrasting sSTOP with cAC-GO trials, we could exclude an primarily attentional role of the rIFG in stopping.

## Methods

### Participants

Sixty-two healthy subjects participated in the study (MEG experiment). The conservative outlier criteria formulated by Congdon et al. (2012) were applied on the behavioral data recorded during the MEG sessions. As a result, one participant had to be excluded because its inhibition stop rate was below 40 %. A second subject had to be excluded because the mean RT of uSTOP trials was higher than the mean RT of cGO trials, indicating that the assumptions underlying the horse race model (Logan et al., 1984) were not fulfilled in this participant. A third subject had to be excluded because no head model could be obtained from the available anatomical MR image.

All of the remaining fifty-nine participants (39 women; mean age, 25 ± 6 years, ranging from 19 to 47) were right-handed according to the Edinburgh Handedness Inventory scale (Oldfield, 1971), had normal or corrected-to-normal vision, and were free of psychotropic medication. None of the participants had a history or current evidence of psychiatric or neurological disorders. All individual participants included in the study were screened for factors contraindicating MEG and MRI scanning and they provided written informed consent before participation. The study was approved by the local ethics committees (Johann Wolfgang Goethe University, Frankfurt, Germany, and Medical Board of Rhineland-Palatinate, Mainz, Germany), and participants were financially compensated for their time.

### Behavioral

Stop signal reaction time (SSRT) was calculated by the mean and integration method. While the integration method tends to underestimate SSRT, the mean method tends to overestimate it (Verbruggen et al., 2013). Therefore we averaged the values of both methods as suggested in Jha et al. (2015). The maximal SSRT (SSRT_max_), as used for the definition of the temporal region of interest (tROI), refers to averaged SSRT values, too.

In the mean method, SSRT was estimated by subtracting the mean SSD from mean RT on go trials. The mean SSD corresponds to the average SSD obtained when using a stair-case procedure that ideally leads to the target of p(respond|signal) = 0.50, which is assumed in the mean method. In the integration method, SSRT was estimated by finding the point at which the integral over the RT distribution equals the actual p(respond|signal) and then subtracting the mean SSD. The integration approximately corresponds to the *n*th RT of the distribution of GO trials (sorted by RT), multiplied by the actual p(respond|signal). For instance, p(respond|signal) is 0.48 across 1000 GO trials acquired, then the *n*th RT is the 480th fastest GO RT. Then SSRT is calculated by subtracting the mean SSD from the 480th RT. The distribution of GO trials also includes incorrect GO trials (Verbruggen et al., 2019).

### Tasks and Procedure

Participants performed a selective stopping task with an additional attentional control condition, i. e. attentional capture go trials, Figure 1). The task and procedure was the same as described in Sebastian et al. (2016). For stimulus presentation, we used Presentation software (version 13.1, www.neurobs.com).

Before the acquisition session, participants had to read the instructions and train the task on a laptop computer for approximately five minutes or until the observed performance confirmed that the task was understood and performed correctly. The acquisition session was split into ten blocks. Throughout the acquisition, participants were asked to hold a response button box with both hands and to respond to the stimuli by pressing a response button with the left or right thumb (MEG) or index finger (fMRI), respectively.

The task comprised three conditions: a GO condition (50 % of all trials), a STOP condition (25 % of all trials), and an attentional capture GO (AC-GO) condition (25 % of all trials). Trials of all conditions began with the presentation of a white fixation cross in the center of the screen with a randomly varied duration between 1,500 and 2,000 ms in the MEG experiment and 500 ms in the fMRI experiment, respectively (Sebastian et al., 2016). Then, a white arrow (GO signal) pointing to the left or right was displayed. Participants were instructed to respond with a left button press to a left pointing arrow and with a right button press for a right pointing arrow. In the GO condition, the white arrow was displayed for 1,000 ms (equivalent to the maximum permitted RT) or until a button press was performed. In the STOP condition, the white arrow was presented first, followed by a change of its color from white to blue after a variable stop-signal delay (SSD). Participants were instructed to try to inhibit their initiated button response after the GO signal. The SSD was adapted to the participants’ performance to yield a probability of 50 % of successful response inhibitions per block. Therefore, a staircase procedure was implemented with the following properties: The initial SSD was set to 210 ms. If the response was not successfully inhibited (unsuccessful stop trial, uSTOP), the SSD in the next STOP trial was decreased by 30 ms with a minimum SSD of 40 ms. If a response was successfully inhibited (successful stop trial, sSTOP), the SSD in the next STOP trial was increased by 30 ms. The maximum SSD was limited by the maximum permitted RT of 1,000 ms. In the AC-GO condition, the white arrow was presented first, followed by a change of its color from white to green after a variable AC-GO signal delay (ACSD), and participants were instructed to continue their response. The ACSD was varied in accordance with the staircase in the STOP condition.

The attribution of color (green/blue) to trial type (STOP/AC-GO) was counterbalanced across participants. In case of an omission error (no button press) in the GO or AC-GO condition, participants were given a short feedback (“oops—no button press”, presented as text for 500 ms) to maintain the participants’ attention and to limit proactive slowing. The length of the intertrial interval (presentation of a blank screen) was 700 ms in the MEG experiment. In the fMRI experiment, the length of the intertrial interval was varied randomly between 2,500 and 3,500 ms. One block consisted of 112 trials presented in a randomized order, resulting in a duration of approximately six minutes (MEG experiment).

### fMRI experiment

#### Anatomical MRI and fMRI data acquisition

A subset of 31 partipicants of the MEG experiment and additional 45 participants performed the task recorded with fMRI (*n* = 76), data are taken from Sebastian et al. (2017). For all participants of the MEG experiment (*n* = 62) an anatomical MRI was recorded. Images were acquired on a Magnetom Trio Syngo 3 T system (Siemens Medical Solutions) at two sites, equipped with an 8-channel head coil at site 1 and a 32-channel head coil at site 2 for signal reception. Stimuli were projected on a screen at the head end of the scanner bore and were viewed with the aid of a mirror mounted on the head coil. Foam padding was used to limit head motion within the coil. A high-resolution T1-weighted anatomical dataset was obtained using a 3D magnetization-prepared rapid acquisition gradient echo sequence for registration purposes (site 1: TR = 2250 ms, TE = 2.6 ms, flip angle = 9°, FOV = 256 mm, 176 sagittal slices, voxel size = 1 × 1 × 1 mm^3^; site 2: TR = 1900 ms, TE = 2.52 ms, flip angle=9°, FOV = 256 mm, 176 sagittal slices, voxel size = 1 × 1 × 1 mm^3^). fMRI images were obtained using T2*-weighted echoplanar imaging sequence (both sites: TR = 2500 ms, TE = 30 ms, flip angle = 90°, FOV = 192 mm, 36 slices, voxel size = 3 × 3 × 3 mm^3^).

#### Preprocessing of fMRI data

SPM12 (www.fil.ion.ucl.ac.uk/spm/software/spm12/) was used to conduct all image preprocessing and statistical analyses, running with Matlab 2013b (MathWorks). Images were screened for motion artifacts before data analysis. Next, images were manually reoriented to the T1 template of SPM. The first five functional images of each run were discarded to allow for equilibrium effects. Then, several preprocessing steps were carried out on the remaining functional images. First, images were realigned to the first image of the first run, using a 6df rigid body transformation. The realigned functional images were coregistered to the individual anatomical T1 image using affine transformations. Subsequently, the anatomical image was spatially normalized (linear and nonlinear transformations) into the reference system of the Montreal Neurological Institute (MNI) reference brain using standard templates, and normalization parameters were applied to all functional images. Finally, the normalized functional data were smoothed with a three-dimensional isotropic Gaussian kernel (8 mm full-width at half-maximum) to enhance signal-to-noise ratio and to allow for residual differences in functional neuroanatomy between subjects.

#### Single-subject analysis

A linear regression model (general linear model) was fitted to the fMRI data of each subject. All events were modeled as stick functions at stimulus onset and convolved with a canonical hemodynamic response function. The model included a high-pass filter with a cutoff period of 128 s to remove drifts or other low-frequency artifacts in the time series. After convolution with a canonical hemodynamic response function, the following three event types were modeled as regressors of interest: correct GO (cGO), successful stop (sSTOP), and correct AC-GO (cAC-GO) trials. Incorrect reactions for each condition and omission error feedback were modeled as regressors of no interest. In addition, the six covariates containing the realignment parameters capturing the participants’ movements during the experiment were included in the model.

#### Group analysis

We compared the neural activation patterns during outright stopping (sSTOP > cAC-GO) by means of a one sample *t*-test. The scanning site was entered as a covariate of no interest. Significant effects for each condition were assessed using *t*-statistics. The results were thresholded at p < 0.05 corrected for multiple comparisons (familywise error (FWE), correction at peak level) and *k* = 5 contiguous voxels. The SPM anatomy toolbox 2.0 (Eickhoff et al., 2007) was used to allocate significant clusters of activation to anatomical regions.

### MEG experiment

#### MEG data acquisition

MEG data acquisition was conducted in line with the good practice guidelines for MEG recordings recommended by Gross et al. (2013). MEG signals were recorded using a whole-head system (Omega 2005; VSM MedTech) with 275 channels. The signals were recorded at a sampling rate of 1200 Hz in a third-order synthetic gradiometer configuration and were filtered with fourth-order Butterworth 300 Hz low-pass and 0.1 Hz high-pass filters. During the acquisition, subject’s head position relative to the MEG sensor array was recorded continuously using three localization coils. To minimize head movements already during data acquisition, we tried to reposition subjects to their individual reference position before a new block if the deviation between the current and their reference position exceeded 5 mm.

For artifact detection of eye movements, the horizontal and vertical EOG was recorded via four electrodes: two were placed distal to the outer canthi of the left and right eye (horizontal eye movements) and the other two were placed above and below the right eye (vertical eye movements and blinks). For heart beat artifact detection, another pair of electrodes were placed on the left and right clavicle (ECG). To ensure a sufficient signal to noise ratio of the recorded artifacts, the impedance of each electrode was kept below 15 kΩ using an electrode impedance meter (Astro-Med).

#### Preprocessing of MEG data

For preprocessing the MEG data and subsequent data analysis, the open source MATLAB toolbox FieldTrip (versions 20120812, 20160202, and 20181010, Oostenveld et al., 2011) was used on the basis of MATLAB (MathWorks, versions 2012b, 2016b, and 2018b, respectively). Trial definition and selective filtering was directed by our aim to identify neural networks related to response inhibition. Specifically, we assumed that the time window in which response inhibition is initiated does not start before the early perceptual processes that are induced by the STOP signal are completed, i. e. 100 ms after STOP signal onset (Amassian et al., 1989). This start point is also motivated by the findings of Swann et al. (2009) and the approach of a pre-registered MEG study (Allen et al., 2018). The definition of the end of the response inhibition time window in sSTOP trials should in principle be based on the SSRT – according to the horse race model that we assumed here. However, we also wanted to directly contrast sSTOP with cAC-GO trials to eliminate attentional processes from the analysis of our selective stopping task. Therefore, we had to take into account that cAC-GO trials might in turn involve button-press related motor activity that is trivially not present in the sSTOP trials. Therefore, the definition of a temporal region of interest (tROI) should preferably end before button-presses in cAC-GO trials set in, but regarding sSTOP trials, the tROI should not end before response inhibition is initiated and finished. To also account for participants with longer SSRTs compared to the average SSRT, we additionally sought to ensure that for these participants the inhibition response process is still captured by the tROI, so we *a-priori* defined a tROI from 100 ms to an SSRT_max_ of 350 ms relative to the STOP signal. To reduce the contamination of our tROI with button-press related motor activity, we rejected cAC-GO trials with an RT_AC-GO_ < SSRT, where RT_AC-GO_ is the “attentional capture GO RT’, defined as duration between AC-GO signal and button press. Median SSRT was 237 ms. More importantly, when contrasting sSTOP and cAC-GO trials, these trials should be also matched in terms of the underlying distribution of go and stop processes according to the independent horse-race model. This is important because the neural correlates of response inhibition revealed by this contrast should not rely on mere timing differences between the underlying go and stop processes (Sánchez-Carmona et al., 2016). Although the distribution of go and stop processes cannot be directly accessed, the horse-race model implies that in sSTOP trials, the response can be successfully inhibited because the go process is slower than the stop process. Thus, to reduce processing speed differences to a minimum, it is necessary to select cAC-GO trials with longer than average RTs, where the underlying distribution consists of slower go processes, comparable with sSTOP trials. Thus, rejecting cAC-GO trials with an RT_AC-GO_ < SSRT additionally supports a close match between the underlying distribution of go and stop processes in both trial types.

Since our analysis focused on the neural correlates of response inhibition, trials were defined with respect to STOP/AC-GO signal onset (Figure 1). Each trial consists of two segments: a task segment of 500 ms duration that started after STOP/AC-GO signal onset, and a baseline segment of the same duration that ended 100 ms before the initial GO signal, while data from intermediate delay periods were discarded for further analysis. If not stated otherwise, only trials with correct behavioral responses were taken into account for data analysis. All analysis steps were based on an equal amount of sSTOP and cAC-GO trials to prevent statistical bias caused by different numbers of trials. Due to the design of the paradigm, the initial number of cAC-GO trials (25 %) exceeded the number of sSTOP trials (approximately 12.5 %). However, RT_AC-GO_ dependent cAC-GO trial selection reduced the difference or even inverted this ratio. When the trial number differed between both conditions for a subject, the minimal amount of trials across both conditions was selected randomly from the condition with more trials.

Trials containing sensor jump or muscle artifacts were rejected using automatic artifact-detection functions provided by FieldTrip. Power line noise was reduced using a discrete fourier transform filter at 50, 100, and 150 Hz. In addition, to remove EOG (horizontal and vertical eye movements) and heart beat related artifacts, independent component analysis (ICA, Makeig et al., 1995) was performed based on trials of all conditions (including GO, STOP and AC-GO), with a duration of 900 ms each, in order to provide a maximal amount of artifacts as input for the ICA. Data was downsampled to 400 Hz before the extended infomax (runica/binica) algorithm was applied to decompose the data (as provided by FieldTrip/EEGLAB). ICs were removed from the data if their spatial topography corresponded with the artifact type (Fatima et al., 2013), which was visually inspected and, when in doubt, correlation coefficients with EOG and ECG data were consulted. Typically, five ICs were rejected (average: 4.9, range: 2 to 8 ICs). Since minimization of head movement is crucial to MEG data quality (Gross et al., 2013), trials were rejected when the head position deviated more than 5 mm from the mean head position over all blocks for each participant.

#### Spectral analysis at sensor level

There are many reasons to favor source-level statistics over statistics at the sensor level; primarily, source level analysis is more sensitive than sensor level analysis (Gross et al., 2013). Therefore, at the first block, the only statistics performed at the sensor level was to define appropriate frequency bands as beamformer parameters for subsequent source reconstruction. We therefore compared the spectral power between 4 and 120 Hz (based on Hanning tapers) over averaged channels during the tROI (100 to 350 ms) with the spectral power of corresponding baseline segments for all participants. For this purpose we used a dependent-samples permutation *t*-test and a cluster-based correction method (Maris and Oostenveld, 2007) to account for multiple comparisons across frequencies. Samples whose *t*-values exceeded a threshold of *α*_cluster_ = 0.05 were considered as candidate members of clusters of spectrally adjacent samples. The sum of *t*-values within every cluster, i. e. the “cluster size”, was calculated as test statistics. These cluster sizes were then tested (two-sided) against the distribution of cluster sizes obtained for 5000 partitions with randomly assigned task and baseline data within each subject. Cluster values below the 2.5th or above the 97.5th percentiles of the distribution of cluster sizes obtained for the permuted datasets were considered significant. Since we aimed to contrast neural power of sSTOP and cAC-GO trials on source level to obtain a set of sources related to response inhibition, we had to use an (orthogonal) statistical test at sensor level in order to avoid circularity (Kriegeskorte et al., 2009). We therefore combined both conditions (sSTOP and cAC-GO trials) for analysis on the sensor level and contrasted these “pooled” task segments with corresponding baseline segments of both conditions.

#### Source Imaging

The anatomical MRI of each participant was linearly transformed to a segmented standard T1 template of the SPM8 toolbox (http://www.fil.ion.ucl.ac.uk/spm/) in MNI space (Collins et al., 1994). This template was overlaid with a regular dipole grid (spacing 1 cm), using an inward shift of –1.0 cm to the brain surface for inside and outside separation. The negative inward shift results in an outward shift that *adds* (not shifts) points to the “inside grid” with a location up to 1.0 cm relative to the brain’s surface, i. e. points outside the brain, that can help to distinguish neural sources from muscle activity. The inverse of the obtained linear transformation was then applied to this dipole grid and the lead field matrix was computed for each of the grid points of the warped grid using a single shell volume conductor model (Nolte, 2003). Since all grid locations of each subject were aligned to the same anatomical brain compartments of the template, corresponding brain locations could be statistically compared over all subjects.

To reconstruct beta and gamma band reactive sources (according to the two frequency clusters revealed by the spectral sensor analysis) a frequency domain beamformer source analysis was performed by using the dynamic imaging of coherent sources (DICS) algorithm (Gross et al., 2001) implemented in the FieldTrip toolbox. Since our experimental setup did not contain any (coherent) external reference, filter coefficients were constrained to be real-valued to restrict our analysis to local source power. This can be done, because we assume that the magnetic fields propagate instantaneously from source to sensor, and therefore, no phase shifts can occur that would lead to complex coefficients (Nunez and Srinivasan, 2006).

Beamformer analysis uses an adaptive spatial filter to estimate the power at every specific (grid) location of the brain. The spatial filter was constructed from the individual lead fields and the cross-spectral density matrix for each subject. Cross-spectral density (CSD) matrices were computed for the task period of 100–373 ms after STOP/AC-GO signal onset and a baseline period of same length, with an offset of –100 ms relative to GO signal onset (see Figure 1 for time scale). To avoid spectral leakage, the length of the time-frequency window should match an integer number of oscillatory cycles of the center frequency of the frequency band width (Harris, 1978). Therefore the length of the tROI and its baseline period was extended accordingly, i. e., ended at 373 ms instead of 350 ms after the STOP/AC-GO signal onset. Frequency bands were defined based on the statistical analysis of spectral power at the sensor level. Thus, CSD matrices were computed in the beta band for 22 Hz (±10 Hz) and the gamma band for 76 Hz (±12 Hz), where spectral smoothing is indicated in brackets. Cross-spectral density matrix calculation was performed using the Field-Trip toolbox with the multitaper method (Percival and Walden, 1993) using four Slepian tapers (Slepian and Pollak, 1978) for the beta band and five Slepian tapers for the gamma band, depending on the required spectral smoothing.

#### Source statistics

To reveal sources related to response inhibition at group level, we statistically tested differences between source power of sSTOP and cAC-GO trials. First, we calculated an activation-versus-baseline *t*-statistic at single-subject level by using an analytic dependent-samples within-trial *t*-test. The *t*-values obtained from this activation-versus-baseline test were then used as input for a cluster-based permutation test for contrasting (“baseline corrected”) sSTOP and cAC-GO trials (dependent samples *t*-test, Monte Carlo estimate, Maris and Oostenveld, 2007). Voxels whose *t*-values exceeded a threshold of *α*_cluster_ = 0.05 were considered as candidate members of clusters of adjacent voxels. The sum of *t*-values within every cluster was calculated as test statistics. These cluster sizes were then tested (two-sided) against the distribution of cluster sizes obtained for 5000 partitions with randomly assigned sSTOP and cAC-GO labels for each subject. Cluster values below the 2.5th or above the 97.5th percentile of the distribution of cluster sizes obtained for the permuted datasets were considered to be significant. In the clusters obtained by our permutation test, we searched for local extrema to identify peak voxels. All of the peak voxels obtained were reported, but for subsequent time-frequency representation (TFR) analysis, only physiologically plausible voxels, i. e., voxels in grey matter according to the Jülich Histological atlas provided by the FSL toolbox (FMRIB’s Software Library, Smith et al., 2004), were used.

### Source time course reconstruction and latency analysis

#### TFRs on virtual channel time courses

To get a picture of the temporal activation pattern of each peak voxel identified by contrasting sSTOP and cAC-GO trials, we analyzed TFRs of these sources. Therefore time courses of these sources were reconstructed as “virtual channels” using bandpass-filtered raw data (2 to 150 Hz). For time course reconstruction, a time-domain beamformer was used (LCMV, linear constrained minimum variance; Van Veen et al., 1997), based on common filters analog to the ones used in the frequency domain. In contrast to the frequency domain, uSTOP trials were additionally included in the common filter computation when a post hoc analysis with uSTOP trials was performed. A principal component analysis on the three reconstructed time courses in *x, y*, and *z* direction was performed for each grid point in order to determine the dominant dipole orientation (direction with the largest variance). The time course of the first principal component was used for subsequent TFR analysis (8 to 44 Hz, Hanning taper with a sliding time window of three cycles were used).

We computed two different TFRs for each source: First, contrast of sSTOP versus cAC-GO trials to reveal inhibition related power changes and second, the difference between sSTOP and cAC-GO trials averaged over all subjects. For both TFRs, as baseline correction, we *z*-transformed sSTOP and cAC-GO trials of each participant separately by subtracting the mean of the corresponding baseline and dividing by the standard deviation of combined (sSTOP and cAC-GO) baseline trials. For the first TFR, representing a group statistics, we used a cluster-based permutation test as described for the spectral analysis at sensor level, except for (time,frequency)-pairs serving as samples. For the second TFR, we subtracted averaged *z*-transformed cAC-GO trials from averaged *z*-transformed sSTOP trials. This absolute difference for each participant and source was used for the onset latency analysis and power correlation.

#### TFR based latency analysis

To answer the question of the temporal order in which the two candidate sources, rIFG and pre-SMA, are activated, we performed an onset latency analysis based on difference TFRs as described above (*z*(sSTOP) – *z*(cAC-GO)). Source power was averaged over the beta band (12 to 32 Hz), smoothed and analyzed for peaks within the tROI. Onset latency (for each participant and source) was defined as the time point at which the averaged source power exceeded a threshold of 25 % of the range between the first positive peak within the tROI and the (positive) minimum found until this peak (in most cases, zero). If there was no positive peak within the tROI for at least one source, we excluded this subject for the TFR based latency analysis. To test for differences between onset latencies of rIFG and pre-SMA at group level, a permutation statistics was performed (50,000 permutations). Onset latency differences were also tested with thresholds of 10 %, 30 %, 50 %, 75 %, and 100 %.

#### Alternative latency analysis using time-resolved SVM

Since we wanted to avoid that putative latency differences were biased due to the way we defined onset latencies on the basis of TFRs, we additionally employed a time-resolved support-vector machine analysis (t-SVM) as alternative. To determine the onset of discriminability in the time-domain between sSTOP and cAC-GO trials, a t-SVM was trained and tested separately for each subject, and for rIFG and pre-SMA. First, we *z*-normalized each source signal relative to baseline period (samples before the GO cue, 500 ms length). Next, to reduce the computational cost and increase the signal-to-noise ratio, the data were smoothed with a Gaussian Kernel of ± 10 ms and downsampled to 300 Hz, similar to the approach in Hebart et al. (2018). After preprocessing, the preprocessed source data were randomly assigned to one of eight *supertrials* (per condition) and averaged (MATLAB code was adapted from Guggenmos et al., 2018). Finally, we separated these *supertrials* into training and testing data with one *supertrial* per condition serving as test data and all the other as training data. The binary classification was performed for each time-point from –100 ms before to 700 ms after STOP or AC-GO signal onset. To obtain a more robust estimate of the classification accuracy, we performed 100 iterations of *supertrials* averaging and classification. The final classification time-series reflects the average across these iterations. For statistical testing, we employed a non-parametric cluster permutation approach (Maris and Oostenveld, 2007), with clustering of subsequent time points. The null-hypothesis of no experimental effect for the MEG classification time series was equal to 50 % chance level and tested from 0 to 350 ms (SSRT_max_).

For each subject we computed the onset and peak latency of the classification time-course. For this, we adopted the method of Marti et al. (2015). First, the data were low-pass filtered at 10 Hz. Secondly, to measure the latency of the peak, we considered all time points, in the window from 0 to 350 ms (SSRT_max_), that exceeded the 95^*th*^ percentile of the distribution of the classification performance. The median of these points was considered as the peak latency. Finally, from the peak latency, the onset of the peak was defined by going backward and identifying the time point at which the classification performance exceeded a certain threshold percentage of the peak. We performed this analysis with different percentage values (10 %, 25 %, 30 %, 50 %) of the difference between the mean decoding accuracy during baseline and the time-window of interest (results were similar across different values). For 13 subjects the decoding performance in the task period did not exceed 2 standard deviations of the baseline accuracy; since also no clear peak was present, we excluded these subjects. Statistical differences between onset latencies of rIFG and pre-SMA at group level were assessed as in the TFR based latency analysis with permutation statistics (50,000 permutations).

### Connectivity

#### Non-Parametric Granger Causality

For the computation of conditional Granger causality (cGC), we employed a multivariate nonparametric spectral matrix factorization (mNPSF). We computed the cross-spectral density matrix of the source signals on the time window from 100 to 350 ms using the fast fourier transform (FFTT) in combination with multitapers (5 Hz smoothing). Using the nonparametric variant of cGC (Dhamala et al., 2008) avoids choosing a multivariate autoregressive model order, which can introduce a bias. Specifically, we used a blockwise approach (Wang et al., 2007), considering the first two PCs of each source signal as a block, and we estimated the cGC that a source 1 exerts over a source 2 conditional on the remaining areas (Bastos et al., 2015). According to our hypothesis, testing for differences between both trial conditions was done for two links, from rIFG to pre-SMA, and vice versa, conditional on all other five active cortical sources as obtained by the source reconstruction. Even though unobserved sources pose an irresolvable problem (Bastos and Schoffelen, 2016), and we can not totally exclude this scenario, we made use of all the information from the sources observed to distinguish between direct and indirect effects and thus avoiding possible spurious results.

#### Statistical testing of cGC

First, we assessed whether the average cGC (in the frequency range of 8–44 Hz) of the source-target pairs (rIFG to pre-SMA and vice versa) was reliably above the bias level, for each condition (sSTOP and cAC-GO) separately. In order to estimate the bias, we randomly permuted the trials 500 times in each condition to create a surrogate distribution of mean cGC values. We tested if the found cGC value was in the upper 97.5 % extreme (equivalent *p* < 0.05 with a Bonferroni correction for two possible source-target pairs) of the surrogates distribution. If the average cGC exceeded the bias level, this source-target link was considered significant. These steps were repeated for each subject separately. Second, for both source-target pairs, cGC values in the sSTOP condition were contrasted with cGC value in the cAC-GO condition at the group level on the subjects that showed at least a significant link in one of the two conditions. The statistical comparison was performed in the range 8–44 Hz using a dependent-samples permutation *t*-metric. A cluster-based correction was used to account for multiple comparisons across frequencies (Maris and Oostenveld, 2007). Adjacent frequency samples with uncorrected *p* values of 0.05 were considered as clusters. Fifty-thousand permutations were performed and the critical *α* value was set at 0.025. A Bonferroni correction was applied to account for multiple comparisons across links.

## Acknowledgements

Parts of this research were conducted using the supercomputer Mogon and advisory services offered by Johannes Gutenberg University Mainz (hpc.uni-mainz. de), which is a member of the AHRP (Alliance for High Performance Computing in Rhineland Palatinate, www.ahrp.info) and the Gauss Alliance e.V. The authors gratefully acknowledge the computing time granted on the supercomputer Mogon at Johannes Gutenberg University Mainz.

## Author contributions

A. S., A. M., P. F., P. J., O. T. designed the experiment. P. J. and M. S. collected MEG data, A. S. and P. J. collected fMRI data. M. S. analyzed the MEG data, E. P. performed SVM and GC analyses, A. S. analyzed fMRI data. M. S., E. P., and O. T. wrote the article. All authors commented on earlier manuscript versions. O. T., M. W., K. L., and P. F. supervised the project.

