## Supplement for "Cortical network mechanisms of response inhibition"

### Supplemental Information

#### Results sensor level

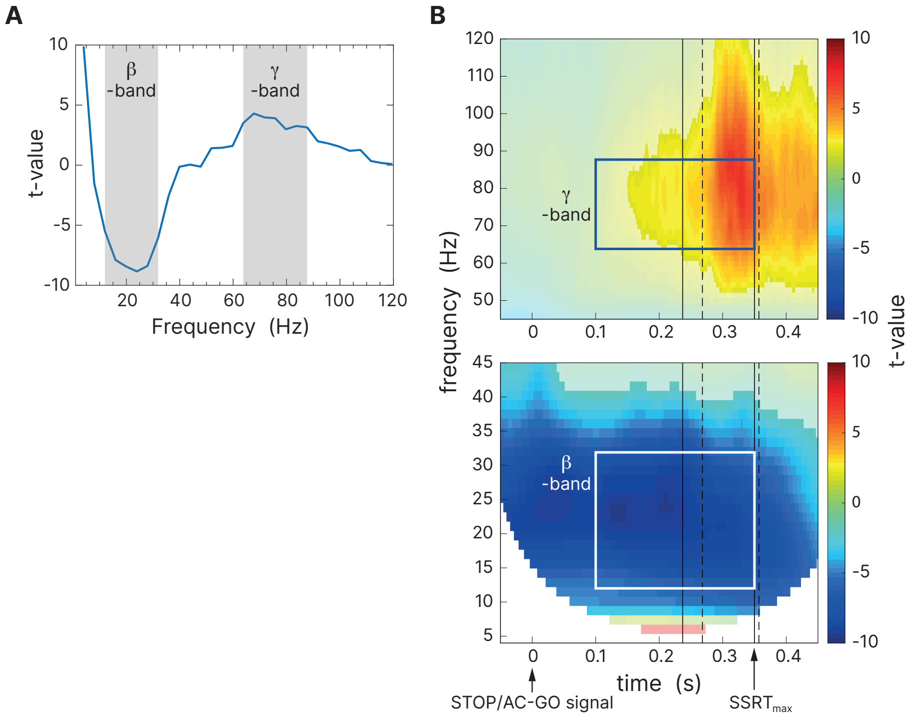


**Figure S1:** Activation-versus-baseline statistics at the sensor level for combined conditions (sSTOP and cAC-GO). Statistics at sensor level were performed in order to find appropriate parameters for source reconstruction. **A**, Two significant frequency bands (highlighted in grey) were revealed by a clustered-based permutation statistics based on spectral power averaged over the temporal region of interest (tROI, 100 to 350 ms) and all sensors (beta band, 12.0 to 31.9 Hz, and gamma band, 63.8 to 87.7 Hz). **B**, Time-frequency representations (TFRs), averaged over all sensors, showed a significant broad-band deactivation in the beta band extending over the whole tROI. Significant activation in the gamma band occured later compared to the beta band. Solid lines represent median and maximum SSRT, dashed lines indicate 10% and 50%-percentiles of $RT_{AC\text{-}GO}$ for selected trials with $RT_{AC\text{-}GO}$ > SSRT. The white/blue box represents the time-frequency window used for soure reconstruction in the beta/gamma band. No significant sources were found for the gamma time-frequency window.

#### Exploratory analysis of uSTOP trials

To clarify if the temporal precedence of rIFG over pre-SMA and the correlation of beta-band power in rIFG with SSRT depend on the outcome of the stopping process (successful vs. unsuccessful stop), we additionally performed the same analysis on uSTOP instead of sSTOP trials. Therefore, the difference between uSTOP and cAC-GO trials was used, instead of the difference between sSTOP and cAC-GO trials.

**Onset latency of beta-band power** When the difference between uSTOP and cAC-GO trials was used for onset latency analysis (see Methods, section *TFR based latency analysis*), the temporal activation pattern did not change, i. e. rIFG was always active before pre-SMA. The permutation test revealed a significant latency onset difference between both sources ($p = 0.0242$, two-sided test, mean onset latency rIFG: $142 \pm 46$ ms, and pre-SMA: $168 \pm 63$ms). This difference refers to a 25 % onset threshold, but it was significant for 10 %, 30 %, 50 %, 75 %, and 100 % thresholds, too. For this test, eight subjects had to be excluded because no positive peak could be found within the tROI ($n = 51$).

**Correlation between beta-band power and SSRT** Beta-band power did not correlate with SSRT when the difference between uSTOP and cAC-GO trials was used (Figure S2), neither for the rIFG ($\rho = -0.062$, $p = 0.336$, Spearman, one-sided test), nor for the pre-SMA ($\rho = -0.094$, $p = 0.740$); two outliers above three standard deviations were excluded from both regressions ($n = 49$).

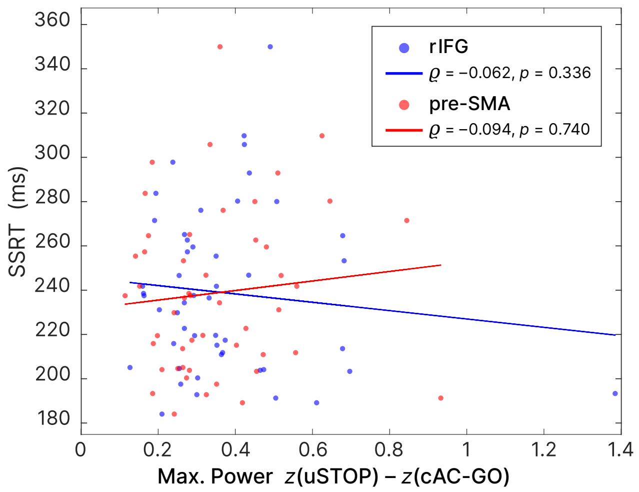


**Figure S2:** The maximum ($z$-transformed) power found in the beta band (12–32 Hz) within the tROI did not correlate with SSRT values, neither for the rIFG, nor for the preSMA when the difference between uSTOP and cAC-GO trials was used.
